# Actin crosslinker competition and sorting drive emergent GUV size-dependent actin network architecture

**DOI:** 10.1101/2020.10.03.322354

**Authors:** Yashar Bashirzadeh, Steven A. Redford, Chatipat Lorpaiboon, Alessandro Groaz, Thomas Litschel, Petra Schwille, Glen M. Hocky, Aaron R. Dinner, Allen P. Liu

**Affiliations:** Department of Mechanical Engineering, University of Michigan, Ann Arbor, Michigan, 48109, USA; James Franck Institute, University of Chicago, Chicago, Illinois, 60637, USA; The graduate program in Biophysical Sciences, University of Chicago, Chicago, Illinois, 60637, USA; Department of Chemistry, University of Chicago, Chicago, Illinois, 60637, USA; Department of Cellular and Molecular Biophysics, Max Planck Institute of Biochemistry, 82152, Martinsried, Germany; Department of Chemistry, New York University, New York, NY, 10016, USA; Department of Biomedical Engineering, University of Michigan, Ann Arbor, Michigan, 48109, USA; Department of Biophysics, University of Michigan, Ann Arbor, Michigan, 48109, USA; Cellular and Molecular Biology Program, University of Michigan, Ann Arbor, Michigan, 48109, USA

**Author notes:** Correspondence: Aaron R. Dinner,; Allen P. Liu. Department of Neuroscience, Baylor College of Medicine, Houston, Texas, 77030, USA. Further information and requests for resources and reagents should be directed to and will be fulfilled by the Lead Contact, Allen P. Liu.

**Keywords:** Actin cytoskeleton reconstitution, actin crosslinker, giant unilamellar vesicles, molecular sorting

## Abstract

How complex actin network architectures arise and coexist in membrane-enclosed cell environment remains unknown. By encapsulating actin and crosslinking proteins α-actinin and fascin in giant unilamellar vesicles (GUVs), we show that physical confinement and its size can lead to formation of complex actin structures, including rings and asters at GUV peripheries and centers. Strikingly, we find that the materials properties of the aster structures depend on the ratio of the relative concentrations of α-actinin and fascin, and we demonstrate that this results from α-actinin and fascin sorting into separate domains in the structures. We complement our experiments with molecular simulations that capture the spontaneous formation of competing network architectures; these provide a microscopic view of the dynamics and delineate the molecular features that promote sorting. We propose that the observed boundary-imposed effect on protein sorting is a general mechanism for creating emergent structures in biopolymer networks with multiple crosslinkers.

## Introduction

The actin cytoskeleton endows cells with remarkable materials properties ^1^. They can resist significant forces but also can deform to migrate through tiny spaces. The spatial organization of the cytoskeleton is critically important for coordinating the forces that enable a cell to move, change shape, and traffic molecules intracellularly ^2–5^. Actin crosslinking proteins crosslink filamentous actin (F-actin) into diverse network architectures, creating complex, heterogeneous materials ^6–8^. α-Actinin is a key actin crosslinker and is enriched in contractile units ^9,10^. α-Actinin dimers are about 35 nm long and bundle F-actin with a spacing that allows myosin binding and in turn contraction. In contrast, fascin is present predominantly at the cell leading edge in sensory protrusions such as filopodia and invadopodia ^11^. There, fascin is found in tight parallel actin bundles with about 6 nm spacing. Biomimetic motility assays with branched actin networks reconstituted on polystyrene beads formed filopodia-like bundles in the shape of aster-like patterns in the presence of fascin ^12,13^.

Live cell and *in vitro* studies have shown that α-actinin and fascin work together in concert to enhance cell stiffness ^14^. In the presence of both α-actinin and fascin, reconstituted branched actin networks on beads formed aster-like patterns and spontaneously segregated into distinct domains: α-actinin was localized to near the surface of the beads while fascin was localized to thin star-like spikes ^15^. Crosslinker size-dependent competitive binding effects of α-actinin and fascin can spontaneously drive their sorting and influence the association of other actin binding proteins ^15,16^. Theoretical models and coarse-grained simulations of two filaments bundling revealed that the energetic cost of bending F-actin to accommodate the different sizes of α-actinin and fascin was sufficient to drive their sorting into domains ^17,18^.

Although biomimetic platforms such as supported lipid bilayers ^19–21^ and the surfaces of giant unilamellar vesicles (GUVs) ^22–24^ introduce appropriate boundary conditions for self-assembly of actin network components into biochemically and mechanically functional networks, they do not confine components in the way that a cell does. To address this issue, actin cytoskeletal components have been encapsulated within or attached to the interiors of lipid-coated single emulsion droplets ^25–27^ or GUVs ^27–30^. Studies that reconstitute actin and microtubule networks from purified components have revealed that spatial confinement can change the structures formed ^31–34^. Here, we investigate the emergent structures and spatial organization of actin networks crosslinked by α-actinin and fascin in a spherically confining GUV environment.

## Results

### α-Actinin induces aggregation and GUV size-dependent formation of actin rings and peripheral asters

To investigate how confinement modulates actin network architecture, we encapsulated α-actinin together with actin inside GUVs of different sizes generated using a modification of continuous droplet interface crossing encapsulation (cDICE) method **(Fig. 1A)**^35^. While no bundling activity was observed in the absence of actin binding proteins **(Fig. 1B)**, α-actinin induced formation of bundled actin filaments in confinement **(Fig. 1C)**. We ‘skeletonized’ *z*-stack confocal image sequences of actin (see Methods and **Supplementary Fig. S1**) to visualize and characterize actin bundles. Actin bundle architecture was found to be highly dependent on GUV size but not α-actinin/actin molar ratio **(Fig. 2A-B)**. Three types of structures were observed: single actin rings near the GUV midplane (**Supplementary Fig. S2**), distinct yet connected actin bundles with no rings (networks), or a combination of connected actin bundles and actin ring(s) around the periphery (ring/network structures). In small (7-12 μm diameter) GUVs, α-actinin-bundled actin merged into a single ring (**Fig. 2C**), similar to structures reported previously ^34,36^, with high probability. The probability of finding single rings was significantly lower in medium (12-16 μm diameter) GUVs, and lower still in large (> 16 μm diameter) GUVs.

**Fig. 1.**
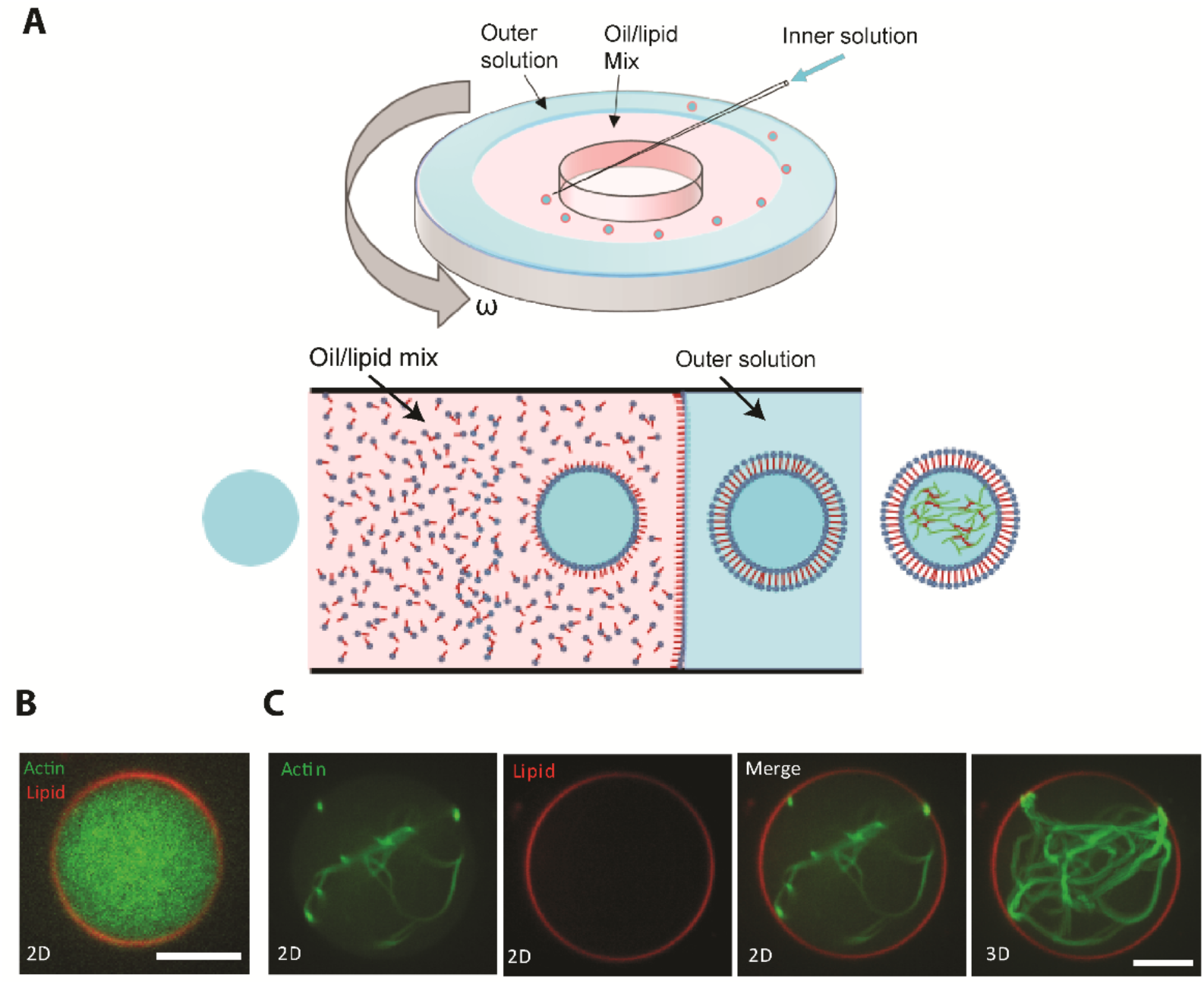
Generation of GUVs with encapsulated actin. **(A)**, Schematic of the setup for generation of GUVs via cDICE. **(B)**, A representative merged (actin:green, lipid:red) confocal image of encapsulated actin in the absence of crosslinkers. Actin, 5 μM, (10% ATTO 488 actin). **(C)**, Representative confocal fluorescence images from an α actinin-crosslinked actin encapsulated into a large GUV. Actin, green (first from left); membrane, red (second from left); merged image (third from left). 3D-reconstructed image from an image stack, right. Actin, 5 μM (10% ATTO 488 actin). α-actinin, 0.5 μM. Scale bar, 10 μm. See also Figure S1.

Increasing GUV size favored more complex actin network structures (**Fig. 2D**) over rings. In particular, in the majority of large GUVs, regardless of α-actinin concentration, F-actin aggregated at the GUV periphery (**Supplementary Fig. S3A, arrows)**, with large clusters proximal to the membrane (**Fig. 2E-F, Supplementary Fig. S3B-C)**. We refer to these structures as ‘peripheral asters’ below; we also describe aster-like structures with clusters in the lumen, and we refer to them as ‘central asters’. We further characterized the peripheral asters by the locations of their bundles and observed that the fraction of GUVs in which all bundles were at the GUV periphery, as opposed to in the lumen, increased with α-actinin concentration (**Fig. 2G)**. The structures formed by α-actinin-actin bundles and their dependence on GUV size are summarized schematically in **Figure 2H**. Given the absence of specific binding between actin bundles and phospholipids, we interpret the preference for the periphery to result from minimization of bundle bending (i.e., elastic energy); the dependence of the bundle location on α-actinin concentration suggests that bundles span the cluster and stiffen as more crosslinkers bind. Stiffening of crossing bundles at the cluster would force actin bundle arms out resulting in the formation asters.

**Fig. 2.**
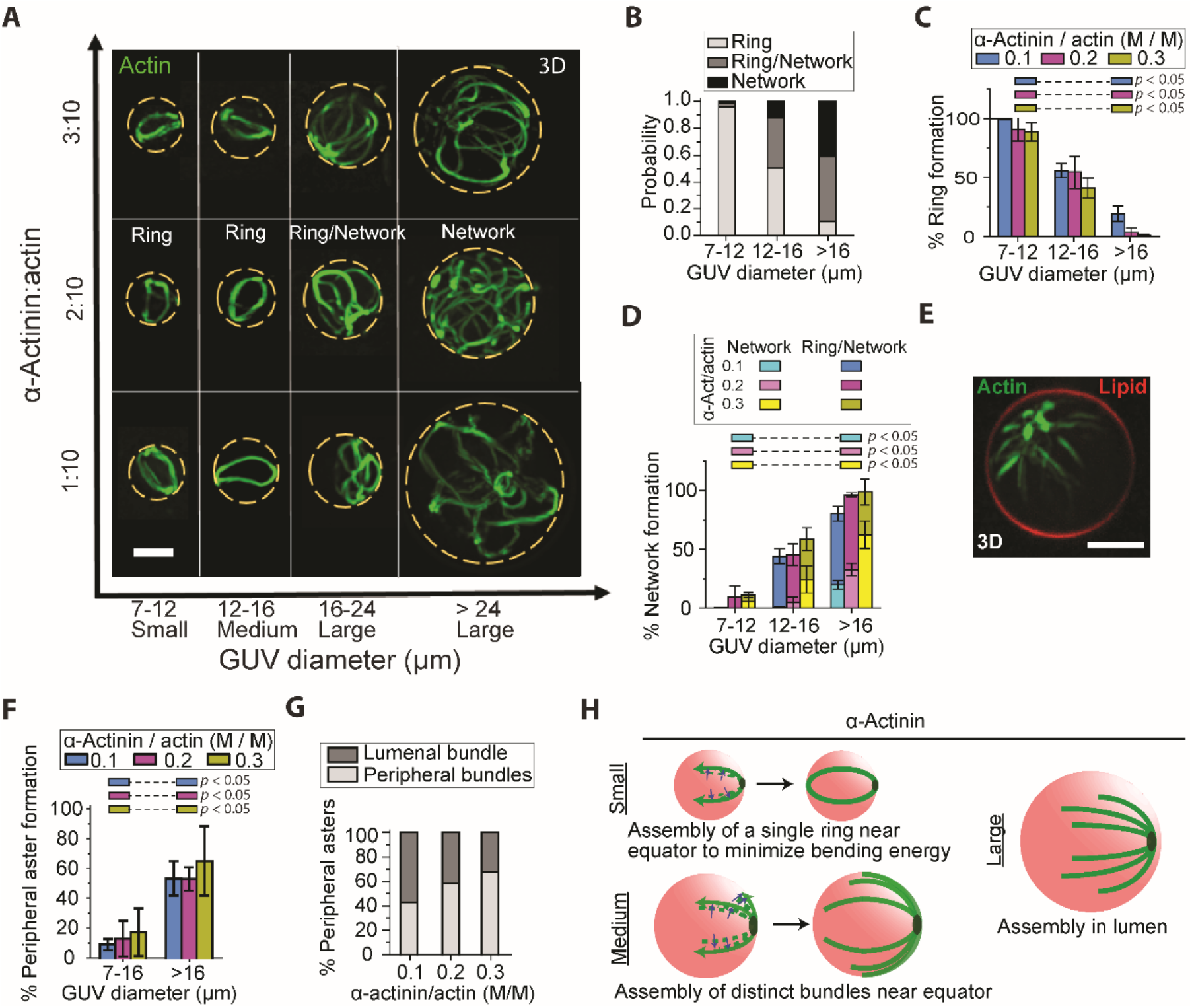
GUV size- and crosslinker concentration-dependent organization of actin-α-actinin networks. (**A**), Representative 3D reconstructed fluorescence confocal images of actin networks of different α-actinin:actin ratios (actin concentration at 5 μM). Actin bundles form networks in larger GUVs, while they form single rings in smaller GUVs. Dotted lines outline GUV boundaries. Scale bar, 10 μm. (**B**), Cumulative probability (for all 3 α-actinin concentrations) of ring and network formation in GUVs with different sizes. (**C** and **D**), Probability of the formation of rings and networks at different α-actinin concentrations. Error bars indicate standard error of the mean; n = 3 experiments. N_GUVs_ per experiment = [389 42 23], [67 45 38], [188 145 128] in order of ascending α-actinin concentration (numbers in brackets are arranged in order of ascending GUV diameter). *p* values compare formation probabilities of rings (C) and networks (D) in small and large GUVs. (**E**), Representative 3D reconstructed image from confocal stack of fluorescence images of an encapsulated α-actinin/actin (3:10 [M/M]) network. High α-actinin concentration can induce the formation of dense clusters at the GUV periphery. Scale bar, 10 μm. (**F**), The probability of the formation of actin peripheral asters depends on GUV size. The majority of actin bundles form peripheral asters in larger GUVs. Error bars indicate standard error of the mean; n = 3 experiments. N_GUVs_ per experiment = [429 23], [112 38], [333 128] in order of ascending α-actinin concentration (numbers in brackets are arranged in order of ascending GUV diameter). *p* values compare peripheral aster formation probabilities for the two ranges of GUV diameters. (**G**), Cumulative (3 experiments) proportion of peripheral asters with peripheral bundles (all actin bundles elongated around GUV periphery) and lumenal bundles (with at least one actin bundle elongated in GUV lumen) in large GUVs (diameter > 16 μm). At high α-actinin concentrations, the majority of actin bundles form asters with all actin bundles elongated around the periphery (peripheral bundles). Number of large GUVs with peripheral asters = [35, 14, 53] in order of ascending α-actinin concentration. (**H**), Schematic summarizing the result of encapsulated actin network assembly by α-actinin (without fascin) in different sized vesicles. Blue arrows show the merging of actin filament bundles (green dashed lines) into distinct peripheral bundles (green solid lines). See also Figures S2 and S3.

In contrast to the results above, the crosslinker fascin formed actin bundles that were sufficiently rigid to stably deform GUVs or were sharply kinked by the membrane as expected ^37^. Fascin alone was unable to induce F-actin clustering or the formation of peripheral or central actin asters.

### Encapsulated α-actinin and fascin together form distinct actin network architectures

The dependence of actin network architecture on GUV size and crosslinker type motivated us to co-encapsulate α-actinin and fascin together with actin in GUVs, and we found that this resulted in the formation of additional actin structures organized around clusters **(Fig. 3A)**. These included central asters, as noted above. Actin clusters at the GUV center were always associated with an aster made up of relatively straight bundles **(Fig. 3A**, yellow arrow). Clusters at the GUV periphery were associated with asters as well, with bundles that curved around the GUV periphery to form partial or complete rings **(Fig. 3A**, white arrows).

**Fig. 3.**
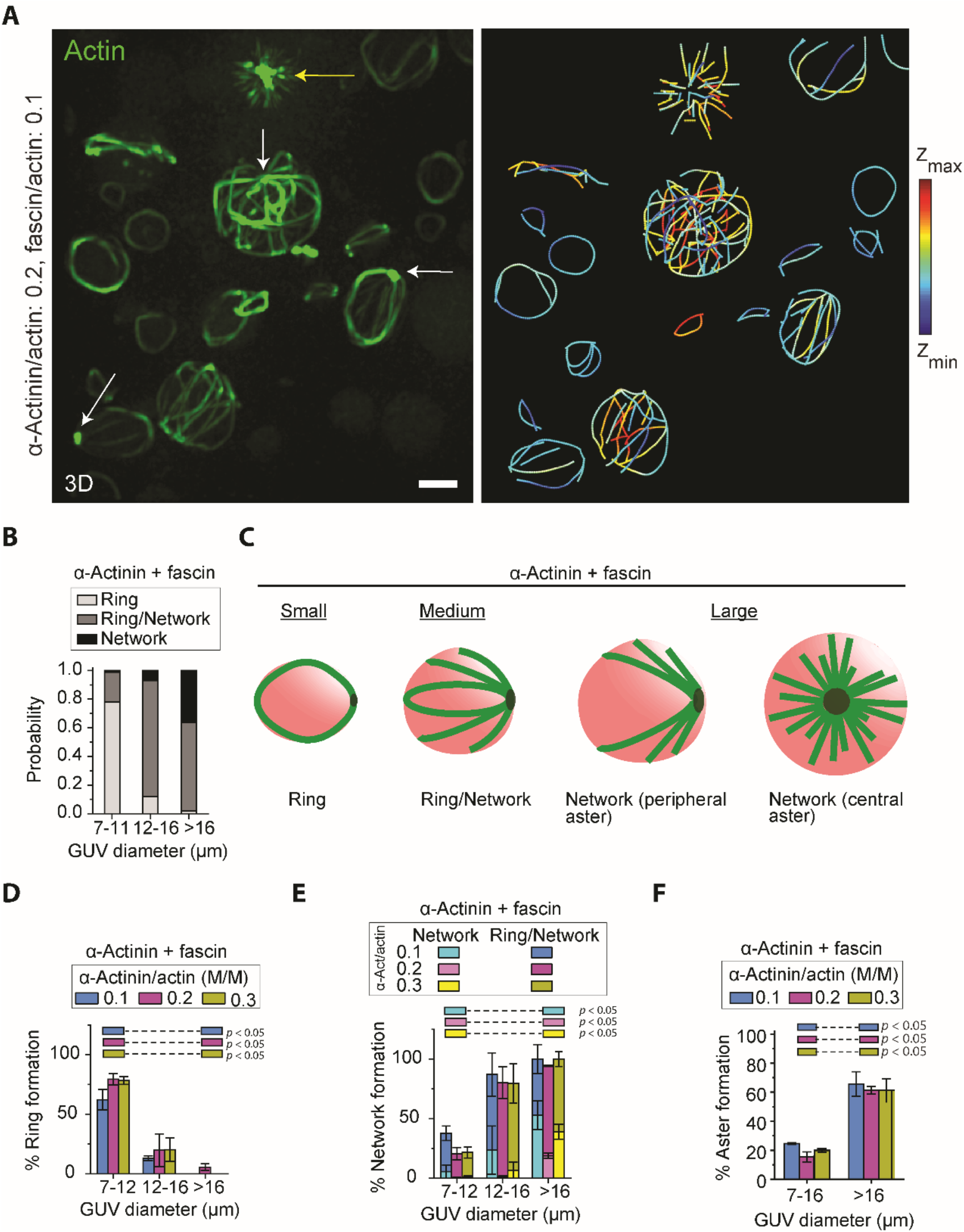
GUV size-dependent formation of rings, peripheral asters, and central asters by α-actinin and fascin. (**A**), Representative 3D-reconstructed (left) and skeletonized (right) images from a confocal fluorescence image stack of 5 μM actin (10% ATTO 488 actin) bundled by 0.5 μM fascin and 1 μM α-actinin in GUVs (composition: 69.9% DOPC, 30% cholesterol, and 0.1% rhodamine-PE). Yellow arrow denotes a cluster of actin fluorescence in a central aster. White arrows denote clusters of actin fluorescence in peripheral asters. Color in the skeletonized image shows z position. Scale bar, 10 μm. (**B**), Cumulative probability (α-actinin concentrations of 0.5, 1, and 1.5 μM with 0.5 μM fascin and 5 μM actin) of ring and network formation in GUVs with different sizes. (**C**), Schematic representation of GUV-size dependent actin networks assembled by α-actinin and fascin. Rings and peripheral asters can occasionally deform GUVs. (**D** and **E**), Probability of the formation of rings (D) and networks (E) at different α-actinin concentrations as a function of GUV diameter. N_GUVs_ per experiment = [101 98 29], [147 58 71], [137 82 24] in order of ascending α-actinin concentration (numbers in brackets are arranged in order of ascending GUV diameter). *p* values compare formation probabilities of rings (D) and networks (E) in small and large GUVs. (**F**), Probability of aster formation (peripheral or central) in the presence of fascin and α-actinin. Aster formation depends on GUV size but not α-actinin concentration. Fascin/actin, 0.1 (M/M). All error bars indicate standard error of the mean; n = 3 experiments. N_GUVs_ per experiment = [199 29], [205 71], [219 24] in order of ascending α-actinin concentration (numbers in brackets are arranged in order of ascending GUV diameter). *p* values compare aster formation probabilities in the two ranges of GUV diameters.

Otherwise, we observed similar trends as previously **(Fig. 3B)**. **Figure 3C** schematizes typical GUV size-dependent actin network architectures formed by α-actinin and fascin. The probability of actin ring formation was reduced dramatically with increasing GUV size (**Fig. 3B-D**), and there was a tendency to form network (and ring/network) structures in medium and large GUVs (**Fig. 3E**). The majority of the network structures in the large GUVs were asters **(Fig. 3F**). All of these trends were insensitive to the α-actinin concentration.

While the probability of aster formation was not observed to be significantly affected by α-actinin concentration, the morphology of asters was. With increasing α-actinin concentration at a fixed concentration of fascin, the cluster in the middle of a central aster grew (**Fig. 4A-B**). Compared to α-actinin-bundled actin structures, the addition of fascin significantly increased the probability of finding central asters **(Fig. 4C)**. In the presence of α-actinin and fascin, increasing GUV size also increased the probability for the cluster to be localized at the GUV center (**Fig. 4D**). The minimization of F-actin bending energy in fascin-associated bundles together with the available space can force filaments to cross close to the center, explaining the tendency for clusters to be in the center of large GUVs. That the size of the central cluster but not the probability of forming a central aster changed with α-actinin concentration implied to us that fascin was driving bundling while α-actinin was driving central clustering. We thus investigated the distribution of the two crosslinkers within an aster.

**Fig. 4.**
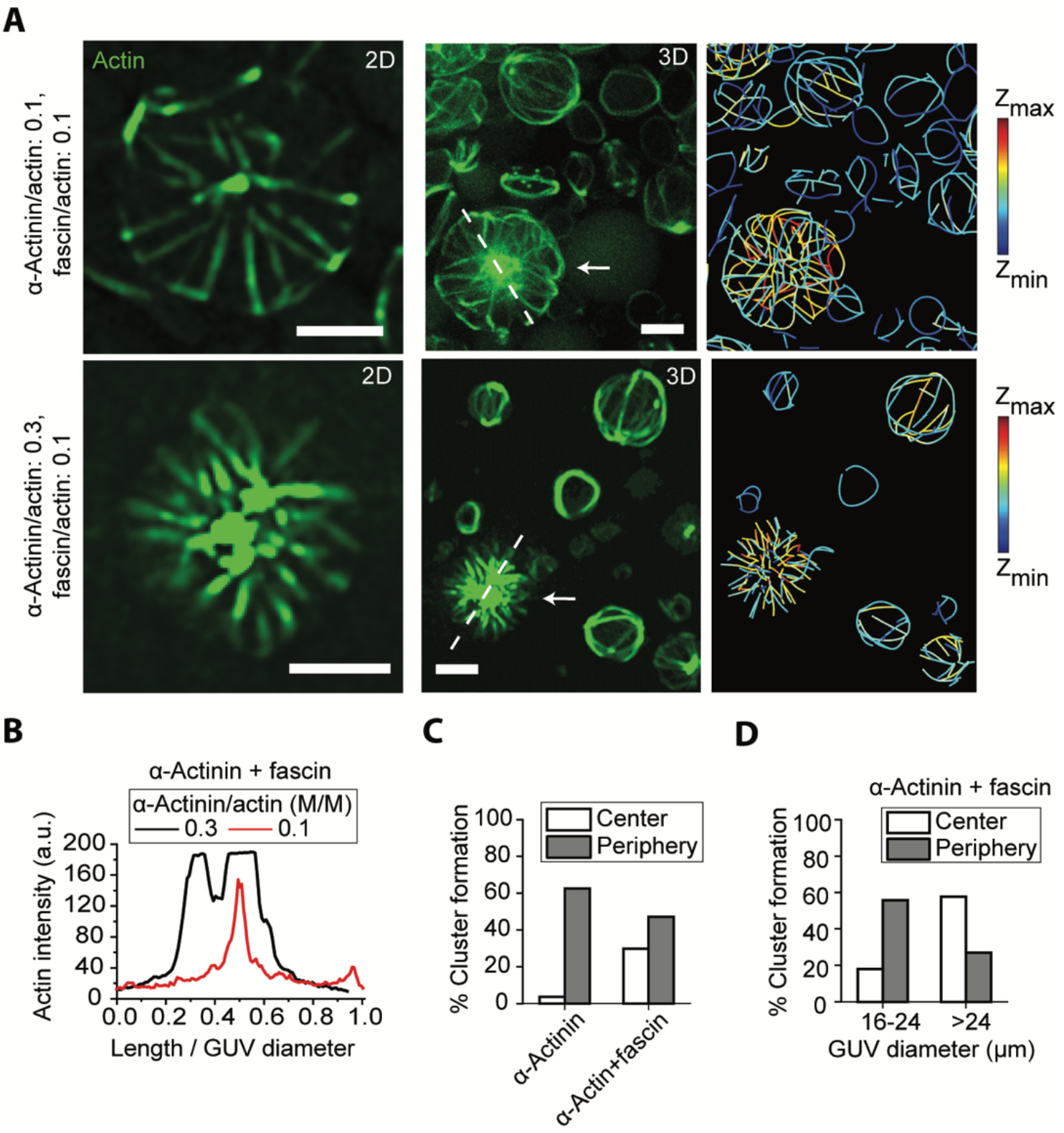
GUV-size dependent localization of F-actin clusters in the presence of α-actinin and fascin suggests crosslinker sorting and domain formation. (**A**), Representative 2D (left) confocal fluorescence images of actin networks shown by arrows (3D reconstructed image, middle) along with skeletonized (right) image of the GUV populations. α-actinin, 0.5 μM (top), 1.5 μM (bottom). Fascin, 0.5 μM. Actin, 5 μM. Scale bar, 10 μm. (**B**), Actin fluorescence intensity along the length of dashed lines drawn across the two GUVs in (A) normalized to the GUV diameters. (**C**), Cumulative probability (for all α-actinin concentrations) of the aggregation of crosslinked actin with/without fascin in GUVs with diameter > 20 μm. α-actinin-fascin-actin bundles tend to shift cluster localization from the periphery to the center of GUVs. Probabilities do not add to 100% because a portion of actin bundles do not cluster at either the center or the periphery. N_GUVs>20 μm_ = 167 [87 (α-actinin), 80 (α-actinin+fascin)]. In our analysis, only a minority of vesicles with encapsulated α-actinin-actin bundle structures and encapsulated α-actinin-fascin-actin bundle structures did not form clusters. (**D**), Cumulative probability (for all α-actinin concentrations) of the aggregation of crosslinked actin in large GUVs in the presence of both α-actinin and fascin. Larger GUVs facilitate centering of clusters. A portion of actin bundles do not cluster either at the center or at the periphery. N_analyzed GUVs>16 μm_ = 87 (61 [17-24 μm], 26 [>24 μm]).

### Crosslinkers spatially sort in central asters

Fluorescently labeled α-actinin was found to localize to clusters in the middle of central asters **(Fig. 5A-C)**. In contrast to central asters, α-actinin was localized entirely along peripheral actin bundles, providing no evidence of spatial segregation **(Fig. 5B, D)**. The absence of α-actinin outside of clusters **(Fig. 5E-F)** supports the hypothesis that the two crosslinkers indeed play very different roles when together. Fascin dominates the region outside the clusters to form tightly packed straight actin bundles in a manner similar to that predicted for Arp2/3 complex-fascin-actin networks in bulk solution ^38^. α-Actinin accumulates in the clusters and crosslinks the rigid bundles, which, in the absence of significant interactions with the membrane, tend to cross at the center to minimize their bending. The size and density of central actin clusters did not change with varying fascin concentration **(Supplementary Fig. S4A-B)**.

**Fig. 5.**
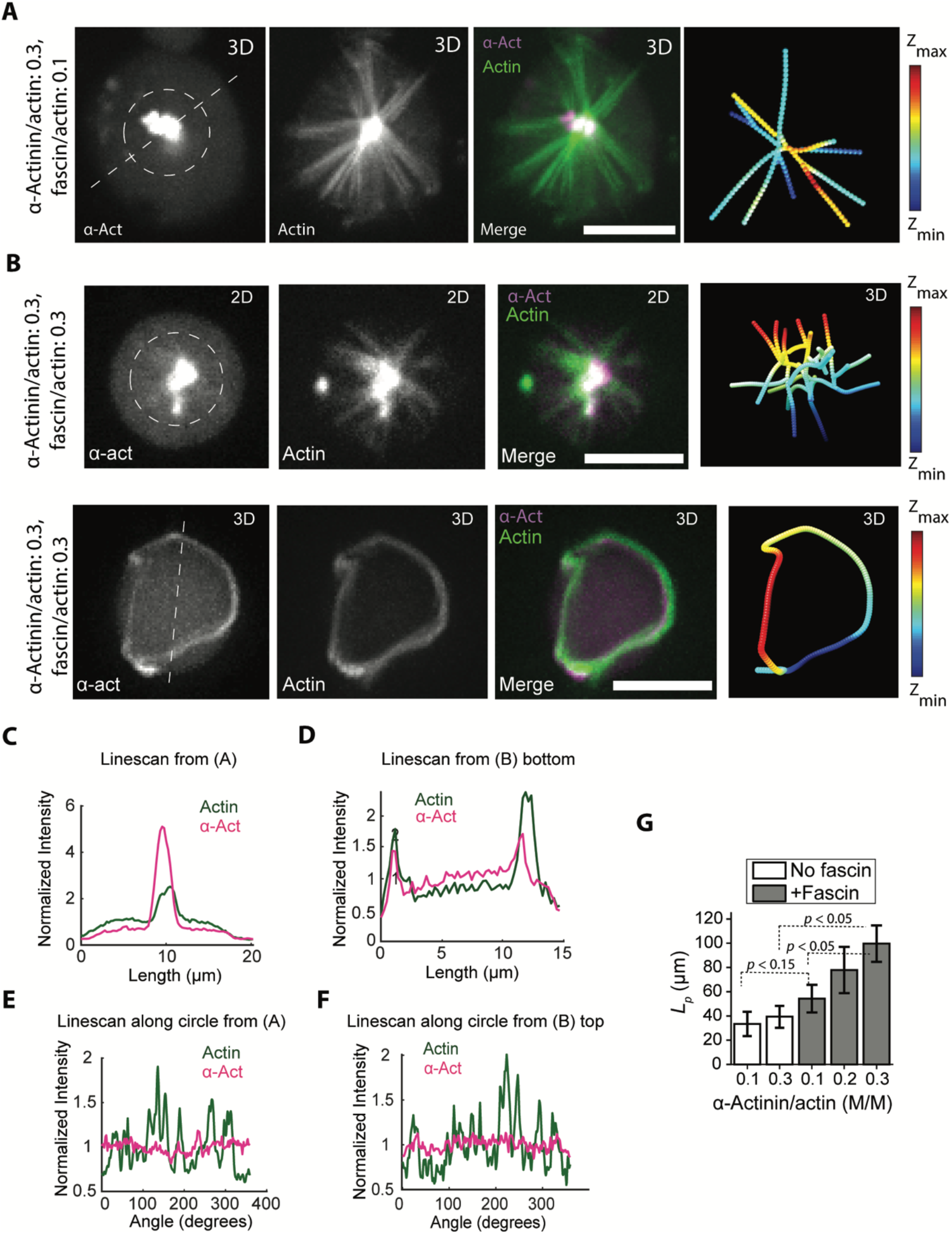
α-actinin and fascin sort and thereby increase bundle flexural rigidity in central aster structures. (**A**), Representative 3D reconstructed confocal fluorescence images of α-actinin, actin, merged, and skeletonized construct of an encapsulated central aster respectively, from left to right. α-actinin, 1.5 μM. Fascin, 0.5 μM. Actin, 5 μM. Scale bar, 10 μm. (**B**), Representative 3D reconstructed confocal fluorescence images of α-actinin, actin, merged, and skeletonized construct of an encapsulated star-like actin network (top) and actin ring (bottom) from left to right respectively. α-actinin, 1.5 μM (top), Fascin, 1.5 μM. Actin, 5 μM. Scale bar, 10 μm. (**C**), Fluorescence intensity of α-actinin and actin along the dashed line drawn across the GUV in (A). (**D**), Fluorescence intensity of α-actinin and actin along the dashed line drawn across the actin ring in (B). (**E**), Fluorescence intensity of α-actinin and actin along the circle drawn around central aster outside the actin cluster in (A). (**F**), Fluorescence intensity of α-actinin and actin along the circle drawn around central aster outside the actin cluster in (B). (**G**), Persistence length of actin bundles without and with fascin (fascin/actin, 0.1 [M/M]) at different α-actinin/actin ratios indicated. *L_p_* [fascin:α-actinin:actin] = 33.3 ± 10 μm [0:1:10], 39.3 ± 9 μm [0:3:10], 54.2 ± 11.4 μm [1:1:10], and [1:1:10], 77.8 ± 19 μm [1:2:10], and 99.7 ± 15 μm [1:3:10]). N_bundles_ = [22 14 17 14 26] in order of *x*-axis categories; 3 GUVs per category. See also Figures S4.

Observing that increasing α-actinin in the presence of fascin increased the size of actin clusters in the GUV center and enhanced the exclusion of α-actinin from actin bundles in central asters, we sought to explore the relation between this exclusion and the rigidity of the actin bundles. The persistence length of actin bundles as a function of α-actinin concentration without and with fascin showed that the bending rigidity of actin bundles is larger in the presence of both crosslinkers compared to α-actinin-bundled actin, as one would expect from the properties of the crosslinkers (**Fig. 5G)**. Strikingly, the persistence length of α-actinin-fascin-actin bundles increased as we increased the molar ratio of α-actinin at a fixed fascin concentration (**Fig. 5G)**. Such an increase in bending rigidity can result from a sorting mechanism in which α-actinin stabilizes a network structure that preferentially recruits further α-actinin to the GUV center and in turn enhances bundling by fascin in the periphery. At low α-actinin concentrations, the sorting is not pronounced, and α-actinin and fascin compete to bundle actin, while, at high α-actinin concentrations, the sorting results in tightly packed fascin-actin bundles with few α-actinin defects. When α-actinin concentration was constant while varying fascin concentration, bundle persistence length **(Supplementary Fig. S4C)** did not change. This suggests that fascin cannot bind F-actin further as fascin-actin bundles outside clusters get saturated.

To understand the microscopic origin of crosslinker sorting in GUVs we turned to coarse-grained simulations of cytoskeletal dynamics. We used the simulation package AFINES ^17,39,40^ and parameterized α-actinin as a relatively long and flexible crosslinker that had no preference for filament orientation and fascin as a short and stiff crosslinker that preferentially bound parallel actin filaments (see Methods). Furthermore, they had different kinetics: fascin had a fast on rate, *k_on_* = 20 *s*^−1^, whereas α-actinin had a slow on rate, *k*_*on*_ = 0.2 *s*^−1^. To compensate for mechanical differences and ensure similar numbers of bound crosslinkers the ratio of *k_on_*/*k_off_* for fascin was 40 while the same ratio for α-actinin was 4. As in the experiments, fascin in the simulations produced tight, well defined bundles, while α-actinin produced flexible bundles that were much more loosely structured **(Supplementary Fig. S5A-D)**. To ascertain whether these physical and kinetic differences can account for the sorting seen in experiment we simulated a 1:1 ratio of the two crosslinkers starting from an initial condition in which actin filaments with fixed lengths crossed close to the center of the simulation region but were otherwise randomly distributed (see Methods). After 100 s of simulation, the α-actinin (**Fig. 6A,** cyan) predominantly occupied the center of the aster, while the fascin (**Fig. 6A,** black) dominated the bundles emanating outwards (**Supplementary Fig. S5E**). To rule out possible model specificity we ran simulations with similar parameters in the package Cytosim ^41^ and saw similar sorting (**Supplementary Fig. S6**).

**Fig. 6.**
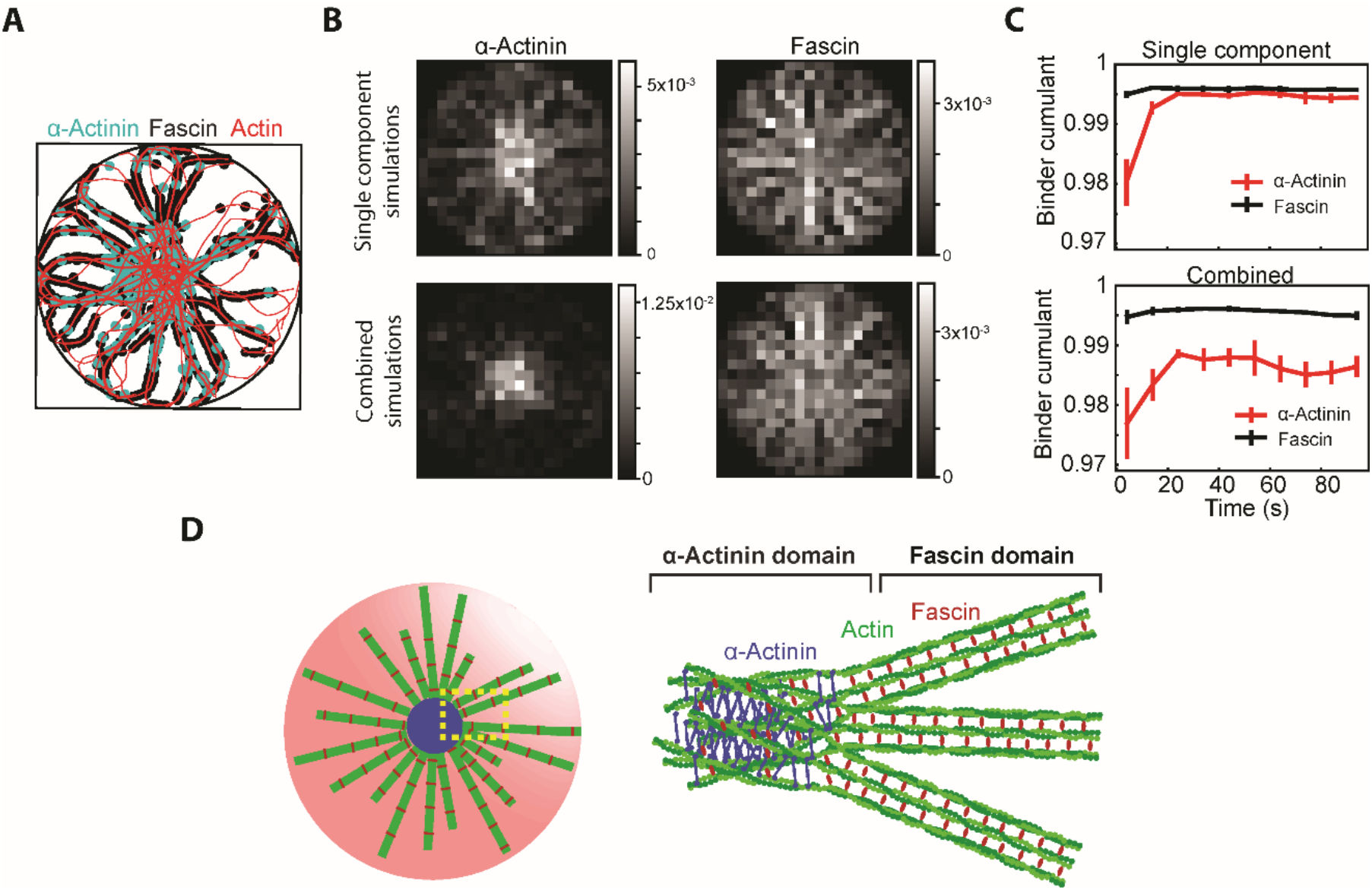
Coarse-grained simulations demonstrate the dynamics of crosslinker sorting in confinement. (**A**), Representative structure after 100 s of simulation of actin filaments (red) crosslinked by α-actinin (cyan) and fascin (black). The border of the containing circle is shown in black. (**B**), PDFs of α-actinin (left) and fascin (right) from either simulation of each alone with actin (top row) or the two in combination with actin (bottom row). PDFs are constructed from the last frames of five independent simulations of 100 s for each condition. (**C**), Binder cumulant as a function of time for each crosslinker separately with actin (top) and the combined simulation (bottom). Values are measured for each simulation independently and then averaged. Error bars represent one standard deviation from the mean. (**D**), Schematic of crosslinker size-dependent sorting in confined α-actinin-fascin crosslinked actin network. See also Figures S5 and S6.

To quantify these observations, we computed the probability density functions (PDFs) of crosslinkers in the simulations. Simulations with a single crosslinker led to relatively flat PDFs (**Fig. 6B**, top). Simulations with the crosslinkers together led to α-actinin PDFs with a steep maximum at the center and fascin PDFs with a shallow minimum at the center (**Fig. 6B**, bottom). To characterize the evolution of the PDFs, we computed the Binder cumulant (see Methods) as a function of time. The Binder cumulant measures the kurtosis of an order parameter, which here is crosslinker density. Higher values correspond to more uniform distributions while lower values correspond to more sharply peaked distributions. The value of the Binder cumulant for the fascin distribution is always high. This indicates that as fascin binds it shows very little spatial preference, forming bundles that span the GUV (**Fig. 6C,** black curves). This is in sharp contrast to α-actinin which, both on its own and in conjunction with fascin, always exhibits a lower value of the Binder cumulant at the beginning of the simulation before relaxing to a steady value (**Fig. 6C,** red curves). This initially low value indicates that, unlike fascin, α-actinin has a propensity to bind in one portion of the aster, i.e., the center, thus resulting in a tighter distribution and a higher value of the Binder cumulant, and only migrates outward into the periphery after these favorable positions have begun to be taken up (**Supplementary Fig. S5E**). While the Binder cumulants in the single crosslinker simulations ultimately reached similar values (**Fig. 6C,** top), when the crosslinkers were together, fascin was able to limit the spread of α-actinin, and a gap between the Binder cumulants persisted (**Fig. 6C,** bottom). These findings confirm that competition-based crosslinker sorting can give rise to the emergent structure observed experimentally.

## Discussion

This work demonstrates that confinement can have a strong effect on reconstituted actin network architectures. We observe rings and both peripheral and central asters, depending on GUV size. As there are no molecular motors in our system, mechanisms that rely on contraction to position the structures ^42–44^ cannot be operative. A recent simulation study showed that confined actin and crosslinkers can form rings, open bundles, irregular loops, and aggregates depending on crosslinker type and concentration, and confinement geometry ^45^. Our results suggest that the positions of crosslinked actin networks can result from minimization of bundle bending as a result of membrane resistance to protrusion. Our experiments and simulations, as well as previous ones ^15,17^, suggest that crosslinker sorting is a factor in determining the architecture of actin bundle networks. Although α-actinin and fascin have similar actin bundling affinities *in vitro*, α-actinin has a higher binding affinity for single filaments and forms parallel and antiparallel bundles with large spacing and mixed polarity as opposed to fascin, which forms tightly packed parallel bundles ^15^. α-Actinin can also form crosslinks between bundles that are neither parallel nor antiparallel ^46^. These properties enable α-actinin to form more complex structures even in bulk, including bundle clusters at high α-actinin concentrations ^14,47,48^. In our system, fascin-bundled actin filaments intersecting and coming in close proximity at the GUV center become crosslinked by α-actinin to form a central cluster (**Fig. 6D)**. This is distinct from the convergent elongation model of how filopodia emerge from a dendritic network ^49^. Similarly, the star-like actin structure with fascin-bundled actin on a bead requires nucleation of branched actin filaments ^12^. Thus, our study reveals a distinct mechanism for creating an emergent actin structure from a two-crosslinker system.

In the sense that the boundary positions filaments to enable this mechanism, one can view it as driving sorting. This may be a general mechanism by which cells can select between different dynamical steady states and tune their functions as a heterogeneous material. Boundary-driven protein sorting can be exploited in a synthetic cell context to generate more diverse self-assembling cytoskeleton structures involving multiple actin crosslinkers. Reconstitution of membrane interfaces has also revealed protein exclusion machineries mediated by protein size and binding energy for effective localization of membrane proteins ^50,51^. Thus, boundary-imposed interactions present a simple yet effective way of protein sorting in cellular processes which induce positioning and activation of proteins at specific sites enabling their localized functions.

## Acknowledgments

We thank Giovanni Cardone of the MPI-B Image Facility for providing FIJI image processing tools. Fluorescently labeled α-actinin was generously provided by the Kovar lab (University of Chicago). We thank Sagardip Majumder and Hossein Moghimian for discussions on experimental procedures and data analysis. APL acknowledges support from National Science Foundation (1612917, 1844132, and 1817909). GMH is supported by a National Institute of Health grant R35-GM138312. ARD acknowledges support from the National Science Foundation (DMR-2011854 and EF 1935260) and the National Institutes of Health (R35GM136381).

## Author Contributions

Experimental design and conceptualization: YB, APL

Experimental methodology: YB, APL, AG, TL, PS

Experimental investigation and analysis: YB

Modeling: ARD, GMH, SAR, CL

Simulations: SAR, CL

Supervision: APL, ARD, GMH

Writing—original draft: YB, APL, SAR, GMH, ARD

Writing—review & editing: YB, APL, ARD, GMH, SAR

## Declaration of Interests

The authors declare no competing interests.

## Methods

### Proteins and Reagents

Actin was either purchased from Cytoskeleton Inc, USA or purified from rabbit skeletal muscle acetone powder (Pel-Freez Biologicals) as previously described ^52^. ATTO 488 actin was purchased from Hypermol Inc, Germany. α-Actinin was purchased from Cytoskeleton Inc. Fascin was either purchased from Hypermol Inc, Germany, or purified from *E. coli* as Glutathione-S-Transferase (GST) fusion protein. For purification, BL21(DE3) *E. coli* cells were transformed with pGEX-4T-3 (GE Healthcare) containing the coding sequences of fascin. Cells were grown at 37 °C while shaking at 220 rpm until the OD_600_ reached 0.5 - 0.6. Protein expression was induced with 0.1 mM IPTG and cell cultures were incubated at 24 °C for 8 h. Cells were harvested by centrifugation at 4,000 x g for 15 min and washed with PBS once. Pellets were stored at −80 °C until the day of purification. Cell pellets were resuspended in lysis buffer (20 mM K-HEPES pH 7.5, 100 mM NaCl, 1 mM EDTA, 1 mM PMSF) and ruptured by sonication. Cell lysates were centrifuged at 75,000 x g for 25 min and supernatants were loaded on a GSTrap FF 1 mL column (GE Healthcare) using an AKTA Start purification system (GE Healthcare) at a flow rate of 1 mL/min. The column was washed with 15 mL washing buffer (20 mM K-HEPES pH 7.5, 100 mM NaCl) and the proteins were eluted with 5 mL elution buffer (washing buffer + 10 mM reduced L-glutathione). Purified products were dialyzed against 1 L PBS twice for 3 h and once overnight at 4 °C. Protein concentration was calculated by UV absorption using predicted molar extinction coefficients (ExPasy) of 110,700 M^−1^cm^−1^. Proteins were concentrated with Centricon filters (Merck-Millipore) when needed and/or diluted to a final concentration of 1 mg/mL in PBS.

### GUV Generation and Microscopy

0.4 mM stock mixture of lipids containing 69.9% 1,2-dioleoyl-sn-glycero-3-phosphocholine (DOPC), 30% cholesterol, and 0.1% 1,2-dioleoyl-sn-glycero-3-phosphoethanolamine-N-(lissamine rhodamine B sulfonyl) (Rhod-PE) in a 4:1 mixture of silicone oil and mineral oil was first made in a glass tube. The lipid/oil mixture could immediately be used or stored at 4 °C for a maximum of 2 days. DOPC, cholesterol, and Rhod-PE were purchased form Avanti Polar Lipids. Silicone oil and mineral oil were purchased from Sigma-Aldrich.

Next, 5 μM Actin (including 10% ATO 488 actin) in polymerization buffer (50 mM KCl, 2 mM MgCl_2_, 0.2 mM CaCl_2_, and 4.2 mM ATP in 15 mM Tris, pH 7.5) and 5% OptiPrep was prepared and kept in ice for 10 min. α-Actinin (0.5-1.5 μM) and/or fascin (0.5-1.5 μM) were then added to the sample. GUVs were generated in 20-30 s after the addition of crosslinkers.

GUVs were generated by a modification of the cDICE method ^35^. A rotor chamber was 3D-printed with Clear resin by using a Form 3 3D printer (Formlabs) and mounted on the motor of a benchtop stir plate and rotated at 1,200 rpm (60 Hz). 0.71 mL aqueous outer solution (200 mM D-glucose matching the osmolarity of inner solution) and around 5 mL of lipid-in-oil dispersion are sequentially transferred into the rotating chamber. The difference in density between the two solutions results in the formation of two distinct layers with a vertical water/oil interface at their boundary. GUV generation was initiated by introduction of 15-20 μL inner solution carrying actin and actin-binding proteins in polymerization buffer containing 5% OptiPrep through a nozzle. Alternatively, a water-in-oil emulsion was first created by vigorously pipetting 15-20 μL of actin polymerization solution (including 5% OptiPrep) in 0.7 mL of lipid/oil mixture and the emulsion was pipetted into the rotating chamber. In either case, generated droplets travel through the lipid dispersion. Lipids are adsorbed at the droplet interface to form a monolayer. As the droplets cross the vertical water/oil interface, they acquire the second leaflet of the bilayer and get released in the outer solution as GUVs.

The outer solution containing GUVs were transferred to a 96 well plate for microscopy. The presence of OptiPrep in GUV lumen increases GUV density and helps GUVs to sediment on the bottom of well plate. Plates were imaged 1 hour after the generation of GUVs unless otherwise mentioned. Images were captured using an oil immersion 60 x/1.4 NA Plan-Apochromat objective on an Olympus IX-81 inverted microscope equipped with a spinning disk confocal (Yokogawa CSU-X1), AOTF-controlled solid-state lasers (Andor Technology), and an iXON3 EMCCD camera (Andor Technology). Image acquisition was controlled by MetaMorph software (Molecular Devices). Actin and lipid fluorescence images were taken with 488 nm laser excitation at exposure time of 350-500 ms and 561 nm laser excitation at exposure time of 20-25 ms respectively. A Semrock 25 nm quad-band band-pass filter was used as the emission filter. Z-stack image sequence of actin and lipids were taken with a step size of 0.5 μm. It should be noted that photo-bleaching of fluorophores significantly impaired actin network self-assembly at the early stages of actin bundling in GUVs. This prevented us from capturing the dynamics of self-assembly by z-stack imaging at a high-temporal resolution.

### Image Processing Methods

Z-stack confocal images of actin-labelled vesicles were mostly analyzed using ImageJ/Fiji ^53,54^, complemented with the plugin Squassh ^55^ from the MOSAIC ToolSuite update site. Encapsulated actin structures were characterized using ImageJ/Fiji in combination with SOAX ^56,57^. Specifically, a skeleton model from the z-stack confocal images was derived as a combination of ImageJ scripts, each one performing a processing step on images. After making the images conform to its input specifications, the software SOAX was launched and controlled through Fiji to determine a model of the filaments in batch mode.

For the cases where GUVs slightly moved during imaging or a drift between z-planes of either encapsulated actin networks or GUVs occurred, images were first aligned by translation using ImageJ plugin StackReg ^58^. The stacks were filtered and the bundles were segmented from them using the Squassh algorithm as implemented in Fiji ^55^. The options for the detection of actin structures with Squassh were as follow: background noise elimination with a window of 1 μm size; regularization factor: 0.075; elimination of background intensity with threshold determined by the Triangle method; Elimination of segmented regions with linear size smaller than 1 μm; automatic estimation of local intensity; Poisson noise model; soft mask for final segmentation. For single GUVs, the image was first cropped in Fiji to include actin bundle-encapsulating GUV of interest prior to image preprocessing (segmentation). Brightness and contrast of all preprocessed images were then manually intensified using Fiji for optimal skeletonization by SOAX program.

A model for actin network on the segmented bundles in each region of interest was determined via SOAX source code ^57^. The program implements a Stretching Open Active Contours method to initiate and generate centerlines, so-called snakes, from each bundle in the population, which is used for quantitative measurements of actin bundles. The SOAX parameter settings to extract the snakes were as follow: Intensity Scaling: 0 (automatic); Gaussian Standard Deviation: 1.2 pixels; Ridge Threshold (Tau): 0.01; Minimum Foreground: 2250; Maximum Foreground: 65535; Snake Point Spacing: 2 pixels; Minimum Snake Length: 17 pixels; Maximum Iterations: 10000; Change Threshold: 0.1 pixels; Check Period: 100; Iterations per press: 100; Alpha: 0.003; Beta: 0.5; Gamma: 2; External Factor: 1; Stretch Factor: 0.3; Number of Background Radial Sectors: 8; Radial Near: 1 pixel; Radial Far: 2 pixels; Background Z/XY Ratio: 3; Delta (snake points): 1; Overlap Threshold: 1 pixel; Grouping Distance Threshold: 2 pixels; Grouping Delta (snake points): 2; Minimum Angle for SOAC Linking: 1.7453 radians; Init z window: marked. Damp z window: unmarked. SOAX generates a file for converged snakes consisting of snake coordinates (actin bundle coordinates). Delete Snake Mode tool of SOAX was used to manually remove converged snakes due to the presence of artefacts or accumulated unbundled fluorescent actin in processed actin images. The SOAX-generated data containing snake coordinates was then saved as a text file. For the cases where the joins between bundles are part of the characterization of actin structures, snakes were cut at junctions and joins formed by SOAX. The coordinates of actin bundles and joints were than save as a text file.

We then developed MATLAB scripts to read the text files, scale voxels to modify image volumes as isotropic, convert pixels to physical units, reconstruct the text as a Chimera marker file in .cmm format for 3D visualization from all directions via UCSF Chimera ^59^ version 1.14, and quantify geometrical features of encapsulate actin bundles.

### Percentages and Probabilities

After taking Z-stack confocal image sequences of GUVs (561 nm) and encapsulated actin (488 nm), and image preprocessing of actin images, 3D reconstructed actin images were obtained via ImageJ brightest point projection using x- and y-axis, separately, as the axis of rotation. Both 3D reconstructed and z-stack images were used for determining the number of GUVs with encapsulated actin bundles including those with certain structural phenotype (single ring, peripheral asters, and central asters). GUV diameters were measured by line scans from both raw actin images and GUVs. GUVs were then categorized as small (7-12 μm), medium (12-16 μm), and large (> 16 μm). The probability of the formation of an actin ring, ring/network, and network per GUV category per experiment were obtained by their count divided by the total number of captured GUVs in the specified category. The percentage of peripheral and central asters in large (or small/medium) GUVs per experiment were obtained by their count divided by the total number of captured large (or small/medium) GUVs with encapsulated actin bundles. The percentage of large (or small/medium) GUVs with actin clusters positioned at the center (or periphery) were calculated by dividing their count by the total number of large (or small/medium) GUVs with encapsulated actin bundles. GUVs encapsulating fluorescent actin monomers with no sign of bundling activity were occasionally found in each population. These GUVs were not taken into account for probability distribution and percentage measurements.

### Calculation of Persistence Length

Using coordinates of the bonds and joints of each skeletonized actin bundle, MATLAB scripts were written to calculate orientational correlation function, <*C*(*s*)> ≡ <cos(*θ*(*s*))>, as a function of arc length *s* along the contour length of selected actin bundles. cos(*θ*(*s*)) is the cosine of the angle between snake bond vectors separated by *s*. < > denotes ensemble average over all snake bonds as starting points. To avoid membrane curvature effect on persistence length measurement, selected actin bundles were among those with no interaction or proximity to the membrane. The lengths of selected actin bundles were 8 < *L* < 20 μm.

Assuming that exponential decay of <*C*(*s*)> in 3D can be described as <*C*(*s*)> = *C_0_*e^−*s*/*Lp′*^, we fitted lines by linear regression to data points (*s*, <Ln(*C*(*s*))>) and determined the slope −1/ *L_p′_* with *L_p′_* denoting the effective persistence length ^60,61^. Among the selected skeletonized bundles, only those with coefficient of determination R^2^ > 0.8 were picked for persistence length measurement. Absolute value of the intercept, Ln(|C_0_|) for selected bundles was found to be around zero with a maximum of 0.03, underscoring the feasibility of assumption C_0_ = 1 for persistence length measurement of actin bundles with length < 20 μm ^62^. We did not find any correlation between persistence length and length of the selected actin bundles for any given experimental condition.

### Statistical Analysis

Error bars in plots of experimental data represent the standard error of the mean. Probability and percentage measurements are based on 3 independent experiments. The reported *p* values are calculated by performing unpaired two-sample student *t*-tests assuming unequal variances.

### Simulation Methods and Analysis

To simulate cytoskeletal networks we used the AFINES simulation package, which is described in detail elsewhere ^17,39,40^ and is summarized here. It employs a coarse-grained representation of components to simulate cytoskeletal dynamics efficiently. Specifically, filaments are modeled as worm-like chains that are represented by beads connected by springs; both the bond lengths and bond angles are subject to harmonic potentials. Similarly, crosslinkers are modeled as linear springs with sites (beads) on each end that can bind and unbind from filaments via a kinetic Monte Carlo scheme that preserves detailed balance. Molecular motors can also be described within this framework but are not included here as they are not present in the experimental system.

The positions of filaments and crosslinkers evolve by overdamped Langevin dynamics in two dimensions. Because the simulation is two-dimensional while the system of interest is not and because we expect the behavior of the experiment to be dominated be network connectivity, we neglect excluded volume interactions between unbound components. However, crosslinker heads can bind to the same filament only if they are greater than a distance *o*_c_ *μm*; apart, where *o*_c_ is a parameter to control the occupancy. While this may lead to quantitative artifacts in the rate of structure formation, previous work has demonstrated that this model is effective in its description of a number of *in vitro* cytoskeletal systems.

In addition to the features detailed in previous work ^17,39^, we have added a circular confinement potential *U*^confine^ to mimic the confinement inside the GUV, and an alignment potential *U*^align^ between actin filaments connected by fascin or α-actinin. The potential energies of filaments (*U_f_*) and of crosslinkers (*U_xl_*) are now

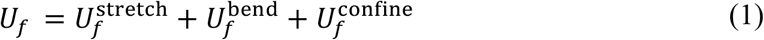

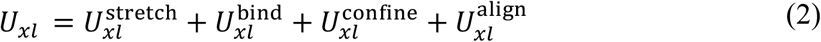

For brevity we only describe the added terms. The confinement potential *U*^confine^ is implemented as a radial confinement potential starting at radius *r_c_* with a tunable force constant *k*_c_,

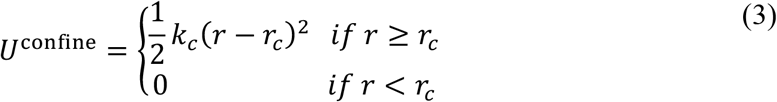

where *r* is the distance of the filament or crosslinker bead from the origin.

For α-actinin we set 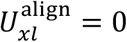. For fascin,

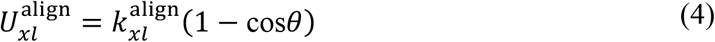

where 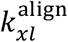 is the penalty parameter for the angle *θ* between the springs of the bound filaments. This term penalizes binding to filaments that are not parallel. The penalty parameter for fascin was set to 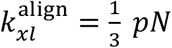 ∙ *μm*;, while the penalty was not implemented for α-actinin.

We utilize these new features and the existing functionality of the AFINES package to model α-actinin and fascin. One chief difference between the two crosslinkers is their respective lengths, with fascin being almost a factor of 6 smaller ^15^. As such, fascin is represented as short, ℓ = 0.1 *μm*;, and stiff, *k* = 1 *pN*/*μm*;, while α-actinin is relatively long, ℓ = 0.5 *μm*;, and soft, *κ* = 0.1 *pN*/*μm*;. Sizes larger than the experimental structures are needed to compensate for the fact that the springs in the simulation are softer than individual proteins, as discussed previously ^17^. That fascin bundles only parallel filaments and α-actinin has no preference between parallel and anti-parallel filaments was captured through the 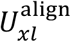 potential, as described above. The simulation parameters are summarized in **Table S1**.

The initial condition of the simulation was generated by first creating each 35-bead filament centered on the origin. After creation, the segment connecting the center to the pointed end was rotated by a random angle drawn uniformly from 0 to 2π. The angle of the other half of the filament was obtained by adding an angle uniformly drawn from between 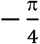 and 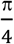 and adding this angle to that of the first segment. Thus, each filament was assigned some random initial bend. Once the angles were assigned, the filament center was moved off of the origin by translating the bead positions in x and y. The amount of displacement in each direction was drawn from a gaussian distribution centered at 0 with a standard deviation of 3 μm. Crosslinker initial positions were drawn with uniform probability from the box centered on the origin with dimensions *r_c_* × *r_c_* where *r_c_* is the radius of the confining circle, which in these simulations is 15 μm. Both crosslinkers are simulated at a density of 5 μm^−2^ within the vesicle. Each simulation ran for 100 s with a time step of 2 ×10^−5^s. The updated version of AFINES including the new features described here as well as the full configuration files used to run these simulations and the script used to generate the initial conditions are available at https://github.com/Chatipat-and-Steven-friendship-forever/AFINES/tree/chatipat_integrate/crosslinker_sorting_in_GUVs. The parameters of the simulation box and actin can be found in **Table S2**.

To characterize the spatial distribution of crosslinkers at a given time, we construct a list of all of the Cartesian coordinates of crosslinkers for which both heads are bound to filaments in that frame. From this list we compute a two-dimensional histogram of crosslinker locations with 20 bins in each dimension. This histogram is normalized by dividing the value of each bin by the area of the bin and the total number of crosslinkers considered. In this manner, we turn our histogram into a probability density function (PDF) that is normalized to one. The images shown in **Fig. 4f** are averages over the PDFs for the final frames of five independent simulations.

We also compute the Binder cumulant, a measure of the kurtosis of the distribution. This measure is frequently used in statistical physics to determine phase transition points in numerical simulations, and is given by

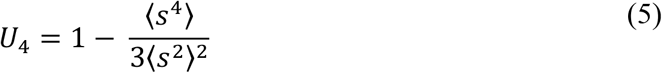

where *s* is an order parameter. In our case, *s* is the value at one point of a crosslinker PDF for an individual simulation, and 〈 〉 denotes a spatial average. The lines shown in **Fig. 4g** are averages over five independent simulations, and the error bars indicate one standard deviation from these averages. The Materials and Methods section should provide sufficient information to allow replication of the results. Begin with a section titled Experimental Design describing the objectives and design of the study as well as prespecified components.

## Notes

### Competing Interest Statement

The authors have declared no competing interest.

