## supplemental figures for "Actin crosslinker competition and sorting drive emergent GUV size-dependent actin network architecture"

### Supplementary Figures and Tables

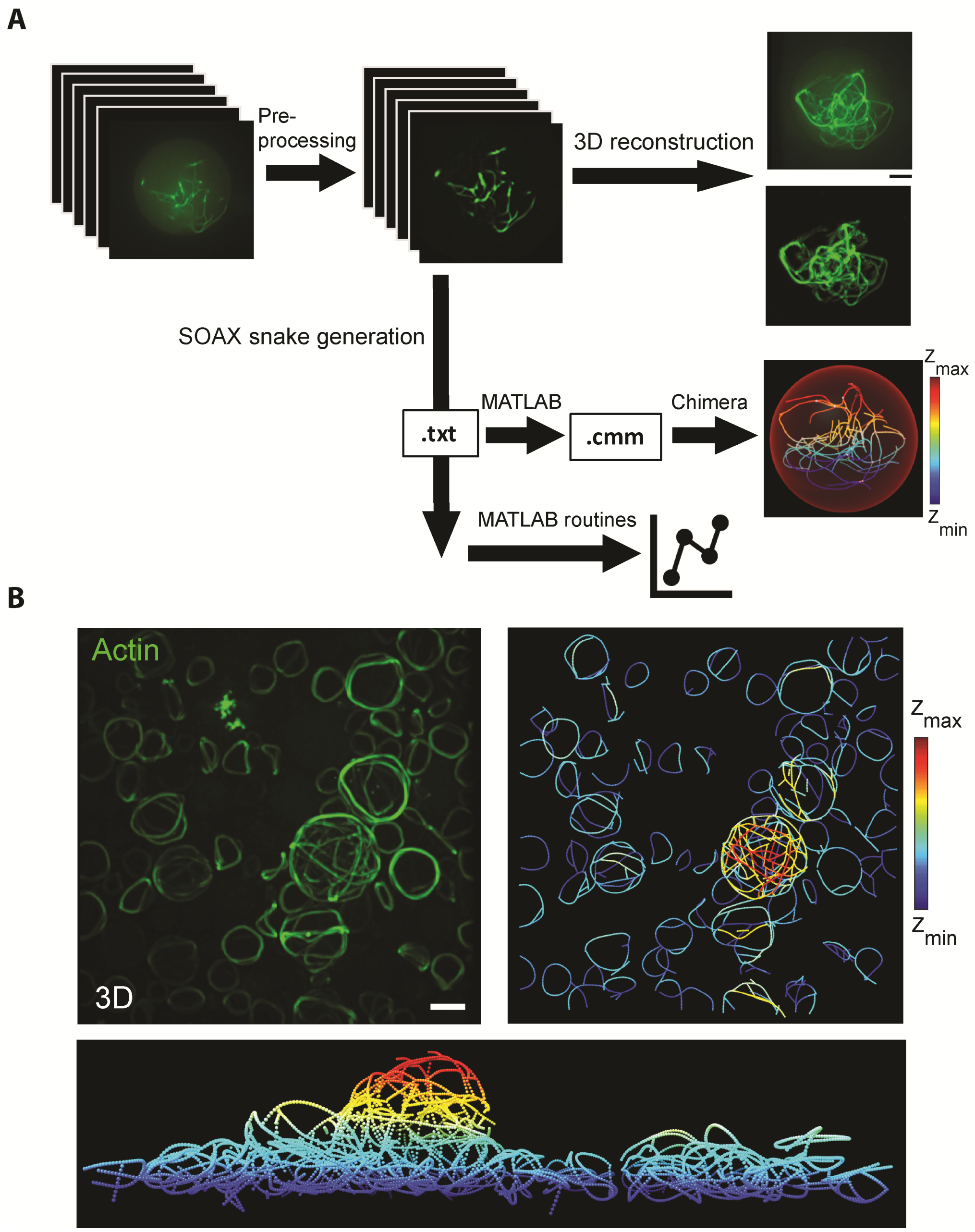


**Fig. S1. Image processing and skeletonization of actin bundles.** (**A**)**,** Flow chart depiction of the steps taken to reconstruct 3D actin images, skeletonize bundles via SOAX software, and MATLAB routines for illustration and characterization of encapsulated actin networks. (**B**)**,** An example illustrating 3D (left), skeletonized (right), and a skeletonized side view (bottom) image of a population of α actinin-fascin-actin (green) networks encapsulated into GUVs with a wide range of sizes. α -actinin/actin, 0.2 (M/M). Fascin/actin, 0.1 (M/M).

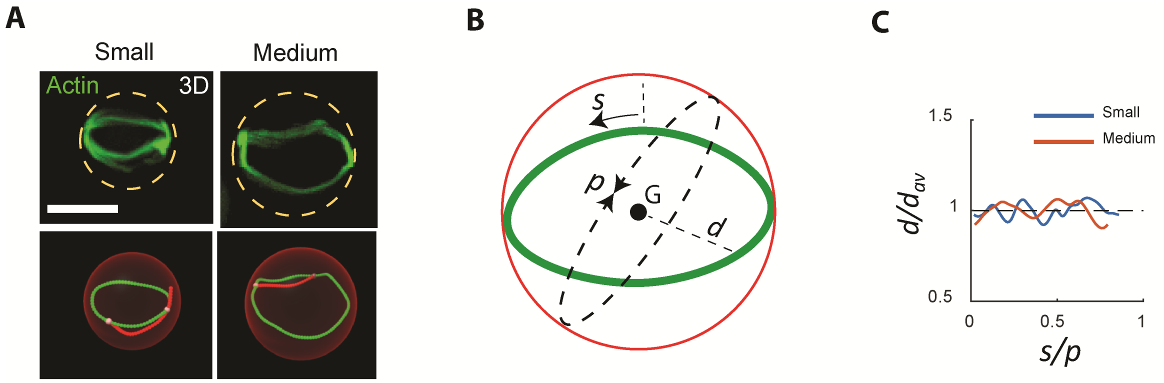


**Fig. S2. Characterization of actin rings.** (**A**)**,** Representative 3D actin images (top) and corresponding skeletonized images (bottom) of actin rings in small and medium GUVs. (**B** and **C**)**,** Slight deviation of an actin ring from a circular geometry due to network formation. Scale bar, 10 µm. (**B**)**,** Geometrical parameters for characterization of actin rings. *d* is the distance between ring center of mass and ring points along the ring in skeletonized structures. $d_{avg}$ is the mean value of *d* (i.e. average ring distance from its center of mass, *G*), *s* is arc length along the ring from a starting point on the ring in skeletonized structures. *p* is the GUV circumference. (**C**)**,** Oscillation over $d/d_{avg} = 1$ depicts the deviation of both rings in (A) (skeletonized images, green) from the locus of a point defining a circle equidistant from ring center.

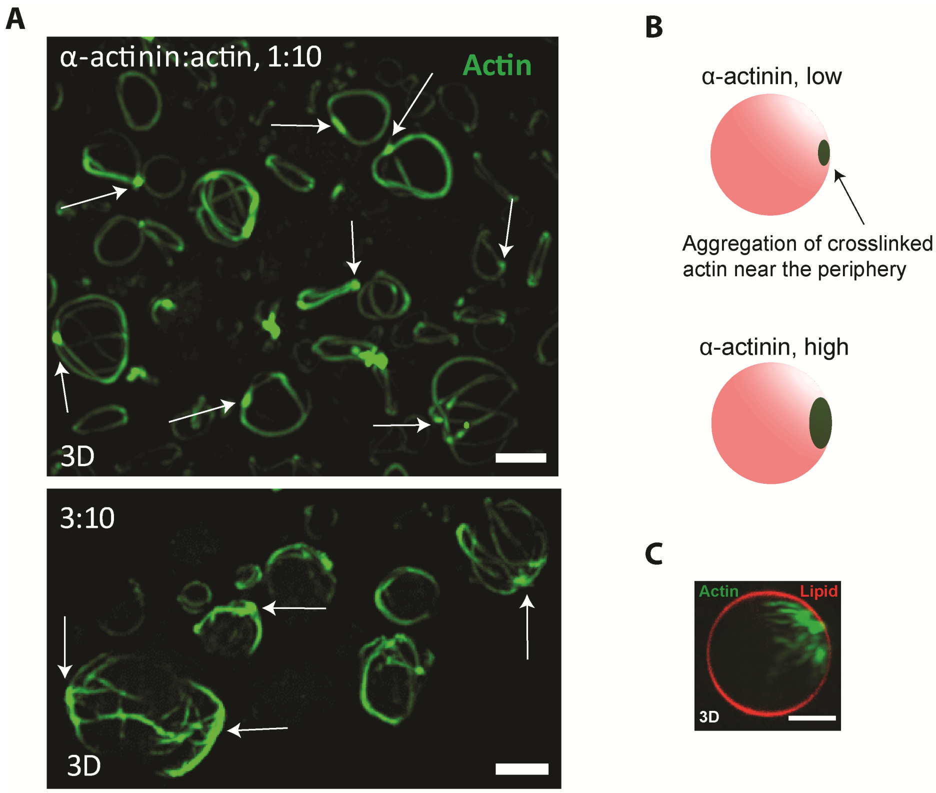


**Fig. S3. α-actinin induces the formation of actin clusters at the periphery of GUVs.** (**A**), Representative 3D reconstructed image from confocal fluorescence images of 5 µM actin with α-actinin at α-actinin to actin ratio of 0.1 (top) and 0.3 (bottom) [M/M]. Crosslinked actin forms aster-like structures emanating from actin aggregates (arrows). (**B**), Schematic representation of actin aggregation (green) to a focal point at the GUV periphery (pink) via cross-linking activity of α-actinin. Higher α-actinin concentration (right) leads to the formation of a larger and highly localized cluster of actin. (**C**)**,** Representative 3D reconstructed image from confocal stack of fluorescence images of an encapsulated α-actinin/actin (3:10 [M/M]) network (rotated view of the same GUV shown in Fig. 1E).

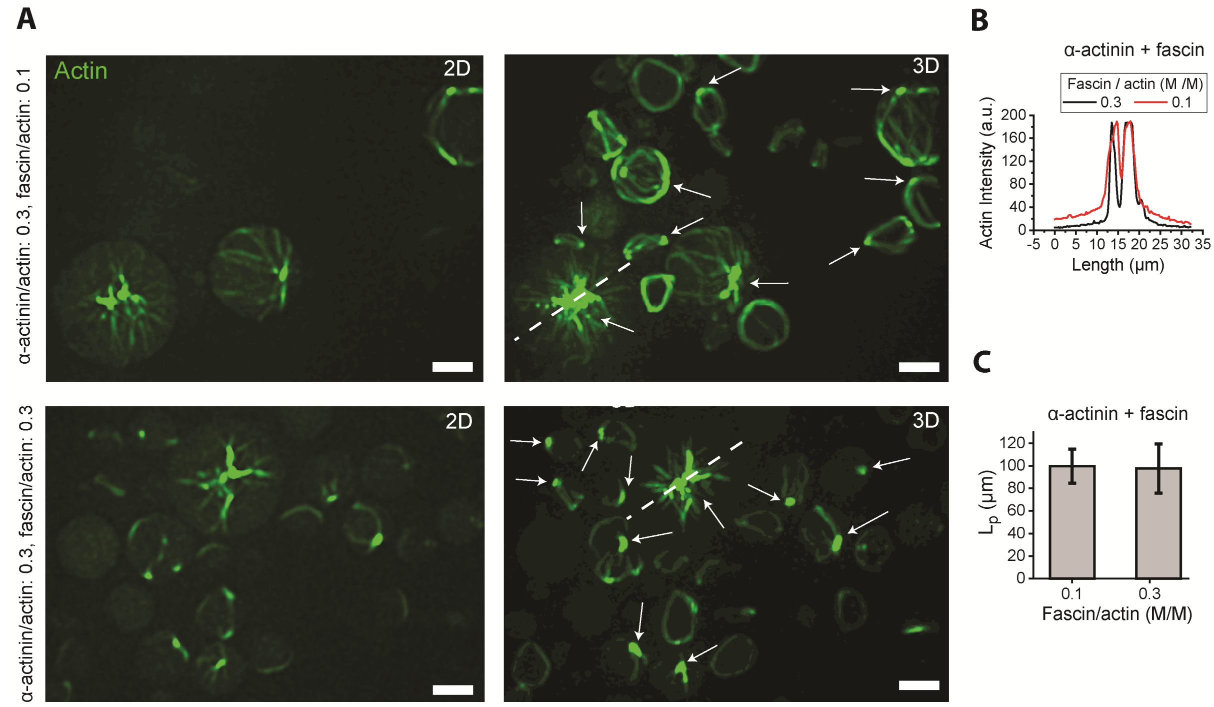


**Fig. S4. At high-α-actinin concentrations, increasing fascin concentration did not change aggregation density nor persistence length of bundles from central asters.** (**A**), Representative 2D (left) confocal fluorescence and 3D reconstructed images of actin networks (right). Actin aggregates are indicated by arrows. Fascin, 0.5 µM (top), 1.5 µM (bottom). α-actinin, 1.5 µM. Actin, 5 µM. Scale bar, 10 µm. (**B**)**,** Actin fluorescence intensity along the dashed lines drawn across the two GUVs in (A). (**C**)**,** Persistence length of actin bundles at two fascin/actin ratios indicated. α-actinin/actin, 0.3 [M/M]). $N_{bundles}$ = [26, 16] in order of x-axis categories; 3 GUVs per category.

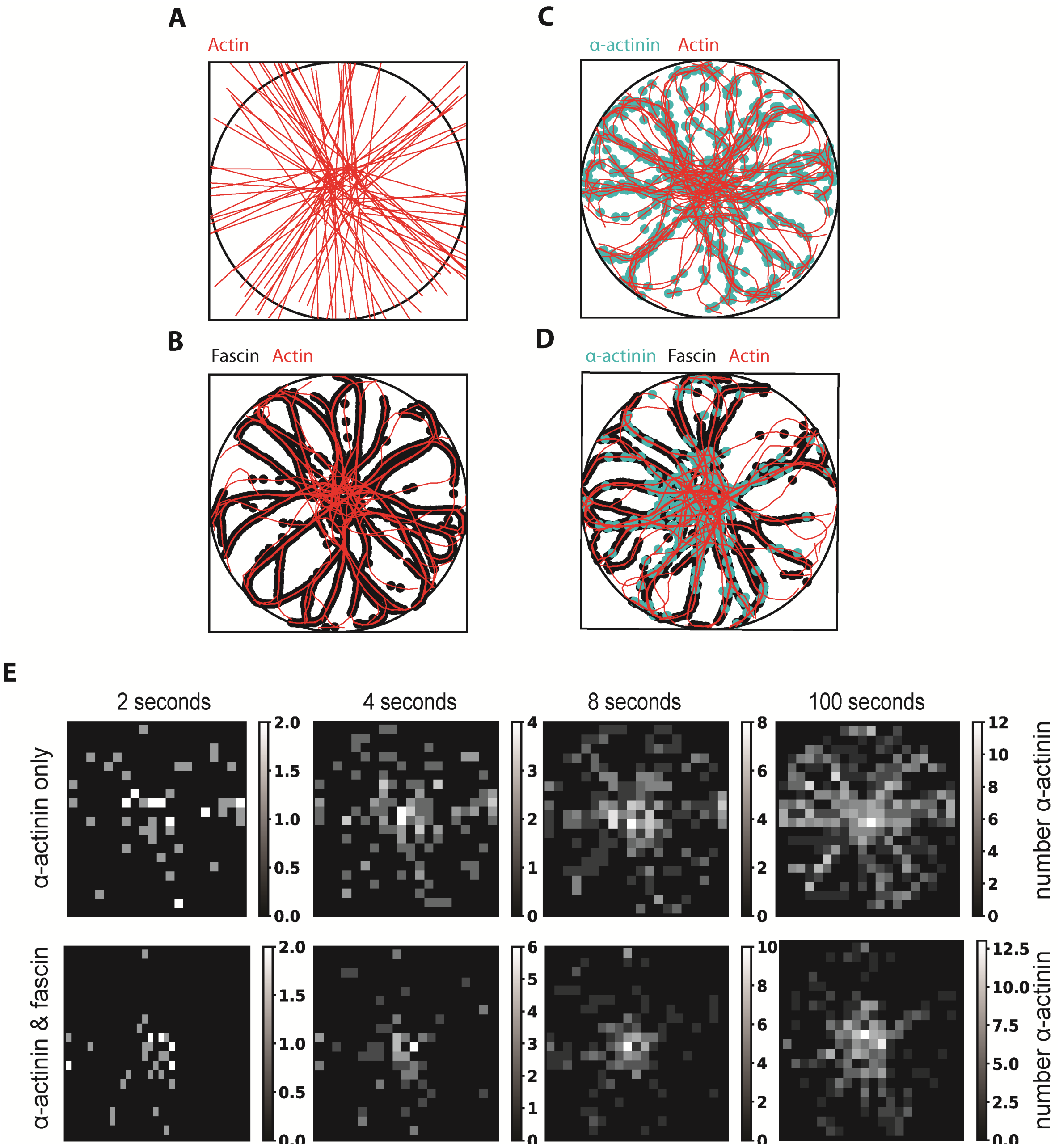


**Fig. S5. Simulated fascin and α-actinin form distinct structures when simulated alone or together on asters.** (**A**), A representative snapshot of the initial condition of one simulation. Actin is shown in red and the boundary of the containing circle is shown as a black outline. (**B**), A representative snapshot from a simulation of fascin alone with actin after 100 seconds of simulation. Fascin is represented as black dots. (**C**), A representative snapshot from a simulation of α-actinin alone with actin after 100 seconds of simulation. α-actinin is represented as cyan dots. (**D**), A representative structure after 100 seconds of both crosslinkers simulated together with actin. **(E),** Time lapse histograms of the number of α-actinin bound in a simulation. The top row is a simulation with α-actinin alone while the bottom row is taken from a combined simulation. Note that in both cases the center of the frame is preferentially occupied first. α-actinin has a propensity for binding in the middle of simulated asters.

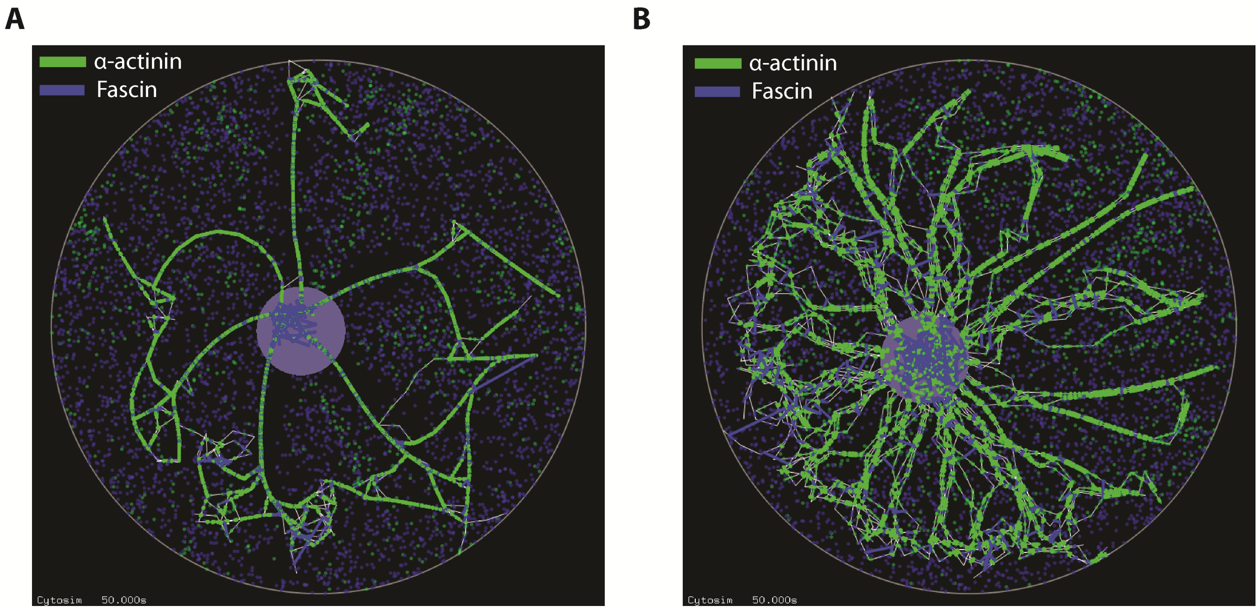


**Fig. S6. Sorting of α-actinin and fascin to distinct regions can be seen by using the Cytosim package.** Structures of asters in Cytosim crosslinked by our representations of α-actinin (blue) and fascin (green) after 50 s of simulation. (**A**)**,** Structure resultant from including fascin's description in the Cytosim fork feature (see Methods). (**B**)**,** Structure resultant from parametrizing fascin not with the fork feature but merely as a crosslink.

| **Crosslinkers properties** | **Fascin** | **α-Actinin** |
|---|---|---|
| resting length ($\mu m$) | 0.1 | 0.5 |
| on rate ($s^{-1}$) | 20 | 0.2 |
| off rate ($s^{-1}$) | 0.5 | 0.05 |
| $k_{xl}^{align}$ $(pN \cdot \mu m)$ | 0.333 | 0 |
| $\kappa$ $(\frac{pN}{\mu m})$ | 1 | 0.1 |

**Table S1.** Simulation parameters of fascin and α-actinin.

| Simulation parameters | |
|---|---|
| X range ($\mu m$) | 100 |
| Y range ($\mu m$) | 100 |
| $r_c$ ($\mu m$) | 15 |
| $k_c$ ($\frac{pN}{\mu m}$) | 20 |
| actin length ($\mu m$) | 0.5 |
| link length ($\mu m$) | 1 |
| polymer bending modulus ($pN \cdot \mu m^2$) | 0.068 |
| number of polymers | 50 |
| beads per polymer | 35 |
| time step (s) | $2 \times 10^{-5}$ |
| end time (s) | 100 |

**Table S2.** Parameters of the simulation box and actin.
